# Altered glutamine metabolism of cultured fibroblasts predicts severity of cardiac dysfunction in the dilated cardiomyopathy with ataxia syndrome (DCMA), a mitochondrial cardiomyopathy

**DOI:** 10.1101/2020.10.11.334938

**Authors:** Melissa A. King, Katherine Heger, Aneal Khan, David Sinasac, Edward L. Huttlin, Steven C. Greenway, Ian A. Lewis

## Abstract

Dilated cardiomyopathy with ataxia (DCMA) syndrome is a rare mitochondrial disorder caused by mutations in the poorly understood *DNAJC19* gene. The clinical presentation of DCMA is very diverse with symptoms ranging from mild cardiac dysfunction to intractable heart failure leading to death in early childhood. Although several lines of evidence indicate that DCMA symptoms are linked to mitochondrial function, the molecular underpinnings of this disease are unclear and there is no way to predict which patients are at risk for developing life-threatening symptoms. To address this we developed a metabolic flux assay for assessing the metabolic function of mitochondria in dermal fibroblasts derived from DCMA patients. Using this approach we discovered that fibroblasts from patients with DCMA showed elevated glutamine uptake, increased glutamate and ammonium secretion, and elevated lactate production when compared to controls. Moreover, the magnitude of these metabolic perturbations was closely correlated with patient cardiac dysfunction. This clinical/metabolic correlation was confirmed in a second blinded cohort of DCMA fibroblasts. Moreover, our metabolic flux diagnostic strategy correctly differentiated severe from mild DCMA cases with only one incorrect patient classification (positive predictive value 1.0 and negative predictive value 0.83). These findings suggest that glutamine catabolism is abnormal in DCMA and may serve as an early biomarker for predicting clinical progression.

**One Sentence Summary:** Alterations in glutamine and lactate metabolism in patient-derived dermal fibroblasts are associated with the severity of cardiomyopathy in DCMA.

## Introduction

Dilated cardiomyopathy with ataxia (DCMA) syndrome is a rare and clinically heterogeneous disorder caused by mutations in the *DNAJC19* gene. DCMA is commonly characterized by deterioration in cardiac function in early life, cardiac conduction delay, unsteady gait, developmental delay, and elevated levels of 3-methylglutaric acid (3-MGC) and 3-methylglutaconic acid (3-MGA) in plasma, urine, and cerebrospinal fluid *(1)*. For reasons that are not well understood, the severity of cardiac disease is highly variable. Some patients present with mild symptoms, such as isolated prolongation of the QT interval, whereas other patients have severe cardiac impairment resulting in end-stage heart failure before two years of age *(2)*. A diagnostic tool predicting clinical outcome for infants born with DCMA could play an important role in management of this disorder allowing early initiation of life-saving cardiac medications.

One major barrier to developing precision medicine approaches for the diagnosis, prognosis, and treatmentof DCMA is our poor understanding of the molecular underpinnings of the disease. Although mutations in *DNAJC19* are known to cause DCMA, how these mutations contribute to clinical phenotype is unclear *(1, 3, 4)*. Recent studies have shown that mitochondria in patient-derived dermal fibroblasts and *in vitro* differentiated cardiomyocytes are highly fragmented, similar to the mitochondria seen in Barth syndrome, a closely related disorder resulting from abnormal cardiolipin remodeling *(5)*. Like DCMA, Barth syndrome results in elevated levels of 3-MGA. However, unlike Barth syndrome, cells from DCMA patients have normal cardiolipin profiles *(6)*. Consequently, DCMA and Barth Syndrome have overlapping phenotypic presentations but appear to differ with respect to the underlying mechanism of disease. In the Hutterite population of southern Alberta, the largest known group of DCMA patients in the world, a common *DNAJC19* mutation is shared amongst all patients and is not predictive of clinical progression. Similarly, levels of 3-MGC and 3-MGA are a clinical hallmark of DCMA, but their levels do not appear to correlate with clinical progression either *(1)*. Interestingly, DCMA patients may also develop hypoglycemia when subject to physiological stress and, in our experience, the administration of glucose can reduce clinical complications. However, glucose levels are also not predictive of disease severity. In summary, many lines of evidence suggest that the clinical presentation of DCMA is linked to mitochondrial metabolism, but the specific molecular mechanisms underpinning the disease remain unclear.

To address this important knowledge gap, we developed a metabolomics-based model system for analyzing mitochondrial metabolic flux in patient-derived fibroblasts *(9)*. Using this approach, we identified abnormal glutamine catabolism in cells from DCMA patients. We show that the magnitude of this phenotype correlates with the severity of cardiac dysfunction. Thus, alterations in glutamine catabolism may provide important predictive information and help enable precision medicine for the management of infants diagnosed with DCMA.

## Results

### DCMA model system using patient-derived fibroblasts

Since the DNAJC19 protein is localized to the mitochondrion and mitochondrial fragmentation is a hallmark of this disease, we predicted that the mitochondrial metabolic function of DCMA patients would be impaired. Moreover, we predicted that metabolic phenotypes observed in well-controlled cell culture systems would correlate with the clinical progression of the disease. To test these hypotheses we studied fibroblasts from six patients (Table 1) representing a transect of clinical cardiac phenotypes ranging from mild (patients with mild or no cardiac dysfunction) to severe (patients with severe dysfunction resulting in death or heart transplantation). Fibroblasts from these patients were cultured *in vitro* and the metabolic activity of these cells was assessed by tracing the catabolism of stable isotope-labelled glucose [U-^13^C] and glutamine [U-^13^C, 2-^15^N]. These nutrients are two of the most important contributors to carbon flux through mitochondria, with glutamine being the preferred carbon source for mitochondrial TCA reactions in fibroblasts *(9)*. Metabolites from these cell lines were analyzed by high-resolution mass spectrometry (MS) and used to quantify metabolic fluxes *(10)*.

**Table 1.** Established fibroblast cell lines and their corresponding phenotype based on patient cardiac dysfunction.

| Cell line name | Phenotype | Age (years) | Sex |
| --- | --- | --- | --- |
| S1 | Severe DCMA | 1.5 | F |
| S2 | Severe DCMA | 1.5 | F |
| S3 | Severe DCMA | 1.5 | F |
| M1 | Mild DCMA | 12 | F |
| M2 | Mild DCMA | 33 | M |
| M3 | Mild DCMA | 35 | M |
| C1 | Control | Adult | unknown |
| C2 | Control | Adult | unknown |
| C3 | Control | 1.5 | M |

Most significantly, we observed alterations in both glucose and glutamine catabolism, the primary carbon inputs for fibroblasts. With respect to glucose metabolism, fibroblasts derived from patients with severe DCMA consumed more glucose (p < 0.005) and secreted significantly more glucose-derived lactate when compared to control lines (1.35-fold, p = 0.0056, Figure 1A). We also observed significantly higher glutamine uptake (1.66-fold, p < 0.0001) and glutamate secretion (2.39-fold, p < 0.0001) in fibroblasts from DCMA patients with both mild and severe cardiac disease in comparison to healthy non-Hutterite control cells (Figure 1B).

**Figure 1.**
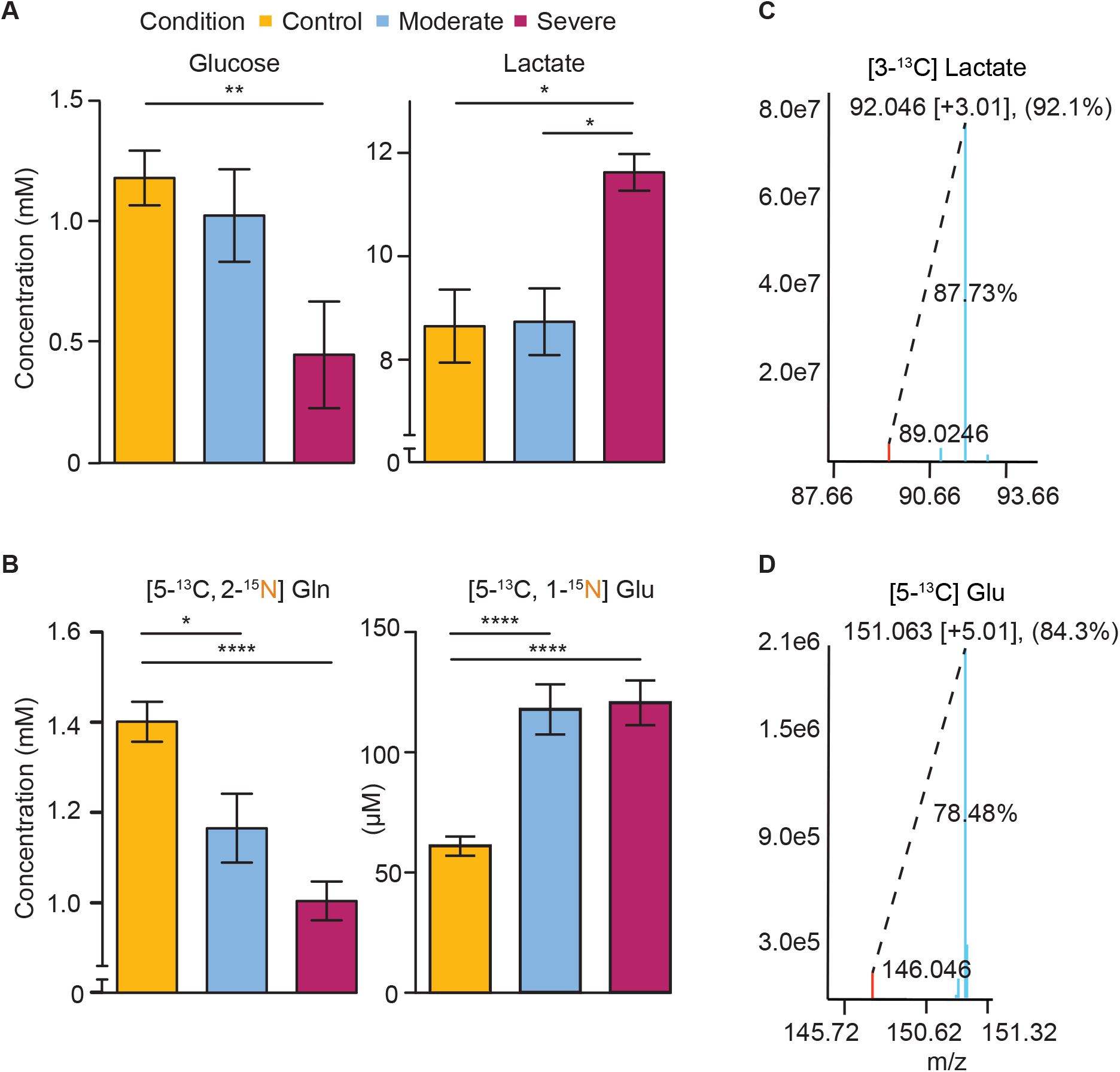
Glucose and glutamine metabolism in control and DCMA fibroblasts. Fibroblasts were grown in DMEM containing either **(A)** U-^13^C-labelled glucose or **(B)** labelled glutamine for 48 hours. Culture medium concentrations reported here represent metabolites observed at the end of the experiment and data are the mean ± SEM of three biological replicates per condition with technical triplicates. * represents p < 0.05 and ** p < 0.005 using unpaired Student’s two-sided t-test (n=3). Mass spectra (right) show **(C)**, the^12^C parent ion for lactate (m/z = 89.0246), and **(D)** glutamate (m/z = 146.046) in red, and the U-^13^C isotopomer for each in blue. Percentages indicate the percent difference in labelling between isotopomers. Abbreviations: Gln, glutamine; Glu, glutamate.

Our isotope labelling studies revealed that in DCMA fibroblasts the lactate was derived almost exclusively from glucose (92.1%) whereas the glutamate observed was produced almost exclusively from glutamine (84.3%). Importantly, these metabolic phenotypes were not attributable to cell numbers as cell lines were normalized by live cell count numbers (Table S1). Moreover, exogenously supplied glutamate, which is not normally found in DMEM, had no impact on these metabolic fluxes. Interestingly, although elevated levels of 3-MGA is a hallmark of DCMA *(11)* we did not detect measurable levels of 3-MGA in the medium of cultured fibroblasts after 48 hours (Figure S1).

### Normal intracellular carbon partitioning

Alterations in glucose and glutamine metabolism could indicate perturbations in the metabolic fluxes of mitochondria from DCMA patients. To assess the impact that DCMA has on intracellular carbon partitioning, we used isotope tracing methods to quantify the relative fluxes through core mitochondrial metabolic pathways *(10)*. Patient-derived DCMA fibroblasts were grown to confluency in ^12^C DMEM. At t=0, the culture medium was replaced with [5-^13^C 2-^15^N] glutamine DMEM and incubated for 72 hours. The medium was then removed and fresh [5-^13^C 2-^15^N] glutamine DMEM was added and incubated for an additional 24 hours before intracellular extraction. This labelling procedure was used to ensure steady-state labelling of TCA metabolites. The pattern of isotopic enrichment in the labelled metabolites was then quantified using high resolution MS and the relative contribution of glutamine versus other carbon sources to intracellular metabolism was analyzed in control, mild, and severe DCMA cell lines.

Surprisingly, the distribution of isotope ratios observed in mitochondrial metabolites associated with the TCA cycle showed no significant differences between DCMA (including patients with both mild and severe cardiac disease) and control cell lines (Figure 2). Of the 35 metabolites measured in our assay, no differences were detected in TCA cycle, urea cycle, or nucleotide biosynthetic intermediates (Figure S2). These data indicate in DCMA fibroblasts the relative contributions of different carbon inputs to central carbon metabolism are stable, even though overall fluxes may be perturbed.

**Figure 2.**
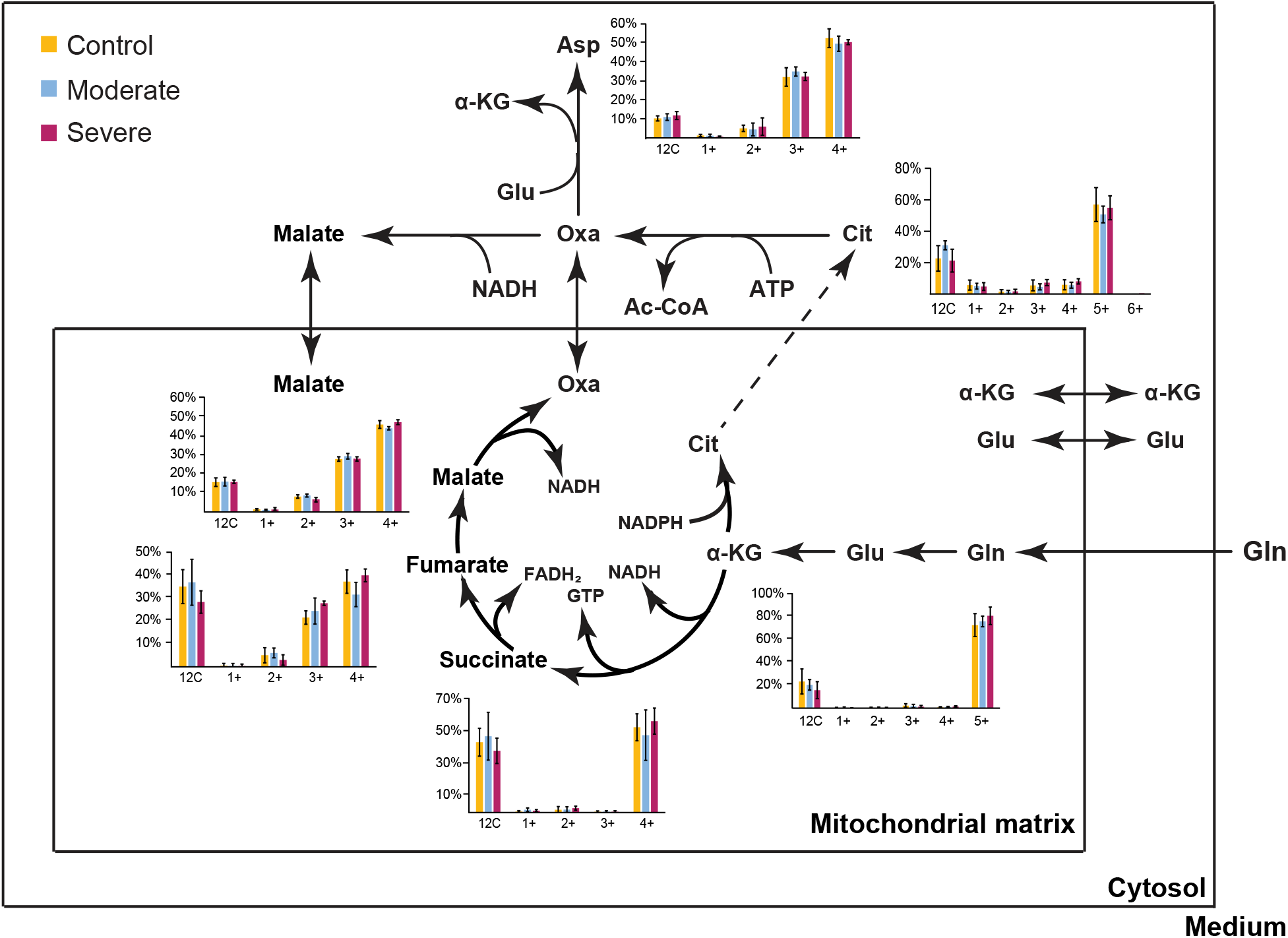
Intracellular carbon isotope enrichment in fibroblasts. Bar graphs show the carbon partitioning percentages of TCA metabolite isotopomers extracted from fibroblast cells lines derived from controls and DCMA patients with mild or severe cardiac dysfunction following incubationwith [5-^13^C 2-^15^N] glutamine for 72 + 24 hours. These data are the mean ± SEM of three biological replicates per condition, with two technical replicates each.

### Altered glutamine utilization in DCMA cell lines

Our data show that DCMA cell lines have elevated glutamine uptake yet maintain relatively normal carbon partitioning between nutrients that feed intracellular metabolic pathways. To better understand the fate of the DCMA-driven glutamine uptake, we conducted a series of isotope labelling experiments to quantify glutamine catabolism and identify compensatory metabolic fluxes that enable this higher uptake without altering downstream carbon partitioning.

Glutamine is metabolized through a series of metabolic conversions that involve both mitochondrial and cytosolic reactions (Figure 3A). It is taken up from the culture medium and then transported into the mitochondrial matrix with a sodium ion *(12)*, where it is catabolized to α-ketoglutarate (α-KG) through the progressive actions of the glutaminase and aminotransferase enzymes. Normally, α-KG is catabolized via the TCA cycle or enters other biosynthetic pathways. However, a certain percentage of this mitochondrial α-KG can be re-converted into glutamine. This process can occur through the exchange with oxaloacetate through the mitochondrial glutamate-aspartate shuttle, and α-KG then enters the cytosol where it undergoes another aminotransferase reaction to produce glutamate. This glutamate can then be converted to glutamine via the energy-dependent glutamine synthetase reaction.

**Figure 3.**
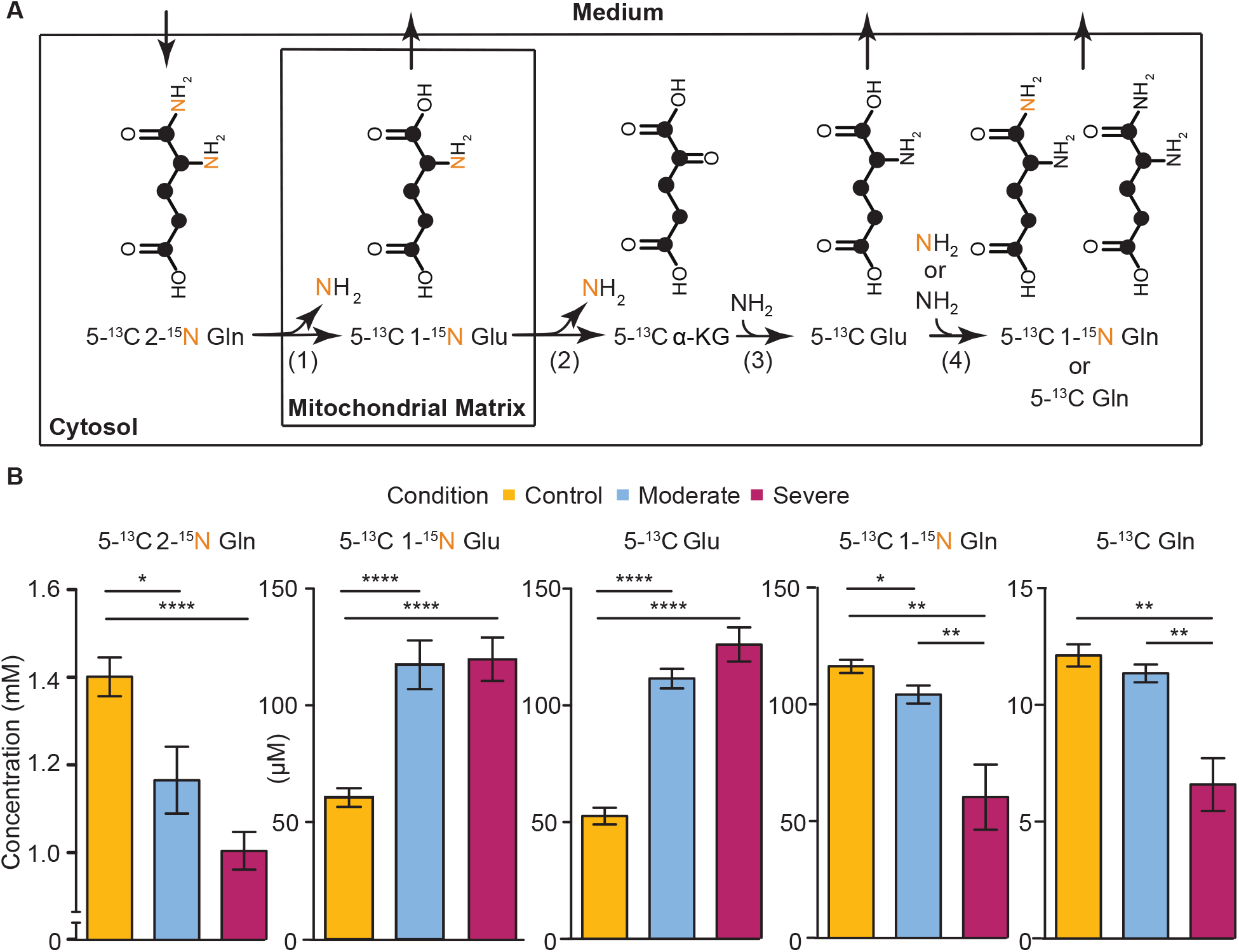
Glutamine metabolism in fibroblasts. **(A)** Labelled nitrogen is represented by a yellow N, and labelled carbon by black dots. Numbers in parentheses represent the enzymatic reactions (1) glutaminase, (2 and 3) aspartate or alanine aminotransferase, (4) glutamine synthetase. **(B)** Products of glutamine metabolism in control, mild, or severe DCMA fibroblasts in medium after 48 hours incubation. Fibroblasts from controls and DCMA patients with mild and severe cardiac dysfunction were provided [5-^13^C 2-^15^N] glutamine in DMEM. These data are the mean ± SEM of three biological replicates per condition with technical triplicates. * represents p < 0.05, ** p < 0.005, **** p < 0.0001 using unpaired Student’s two-sided t-test (n=3). Arrows indicate net flux observed in cultured fibroblasts. Abbreviations: Gln, glutamine; Glu, glutamate.

This energy-consuming recycling of glutamine can be monitored by following the isotope labelling patterns of [5-^13^C 2-^15^N] glutamine (Figure 3A). To better understand the fate of glutamine in DCMA cell lines, we used high resolution MS to follow these isotopomers and quantify the catabolism and re-synthesis of glutamine. Using this approach, we identified significant changes in glutamine recycling that correlated with clinical severity (Figure 3B). As shown previously, double-labelled glutamine levels in the medium were significantly lower in DCMA cell lines. Moreover, [5-^13^C ±^15^N] glutamate levels were significantly elevated, suggesting a net deamination of glutamine. In addition, [5-^13^C 1-^15^N] glutamine levels, which are indicative of the synthesis of glutamine from fully labelled glutamate, were significantly lower in DCMA cell lines (p = 0.0079 and 0.0011, 0.52- and 0.54-fold in severe compared to control and mild DCMA cell lines respectively). This pattern was also seen with [5-^13^C] glutamine levels, which are the product of glutamine synthesis from labelled glutamate. Overall, these data suggest that in DCMA cell lines there is a net uptake of glutamine that is commensurate with a deamination of glutamine and glutamate secretion. Moreover, this process appears to repress glutamine recycling.

### Altered nitrogen secretion in DCMA cell lines

Our isotope labelling data showed increased uptake of glutamine that was associated with increased secretion of labelled glutamate. This net reaction results in a loss of nitrogen which has multiple possible fates within the cell. One possibility is that this nitrogen is lost through the mitochondrial glutaminase reaction, which would result in the net production of ammonium. Although this ion normally enters the urea cycle, our intracellular labelling data showed no alterations in urea cycle labelling, suggesting an alternative excretion route for glutamine-derived ammonium. Ammonium can also be secreted directly into the medium. To quantify nitrogen loss through this direct secretion mechanism, we used ^1^H-^15^N heteronuclear single quantum coherence (HSQC) nuclear magnetic resonance (NMR) spectroscopy to quantify glutamine-derived ammonium levels in the media of cells incubated for 24 hours in the presence of 2-^15^N glutamine. Ammonium levels were quantified using established methods *(13)* and data were analyzed using rNMR *(14)*.

We observed ^15^N-labelled ammonium in both the DCMA and control cell lines, with the severe DCMA cell lines showing significant (p = 0.0027, 1.23 fold) elevation in ammonium secretion (Figure 4). These data indicate that the compensatory [5-^13^C] glutamate secretion observed in DCMA cell lines is balanced by direct nitrogen secretion to the medium. DCMA-linked perturbations in glutamate secretion and ammonium production were both ∼100 µM higher than control lines (105 µM glutamate and 106 µM ammonium) suggesting that all of the altered glutamate secretion could be attributable to the glutaminase reaction.

**Figure 4.**
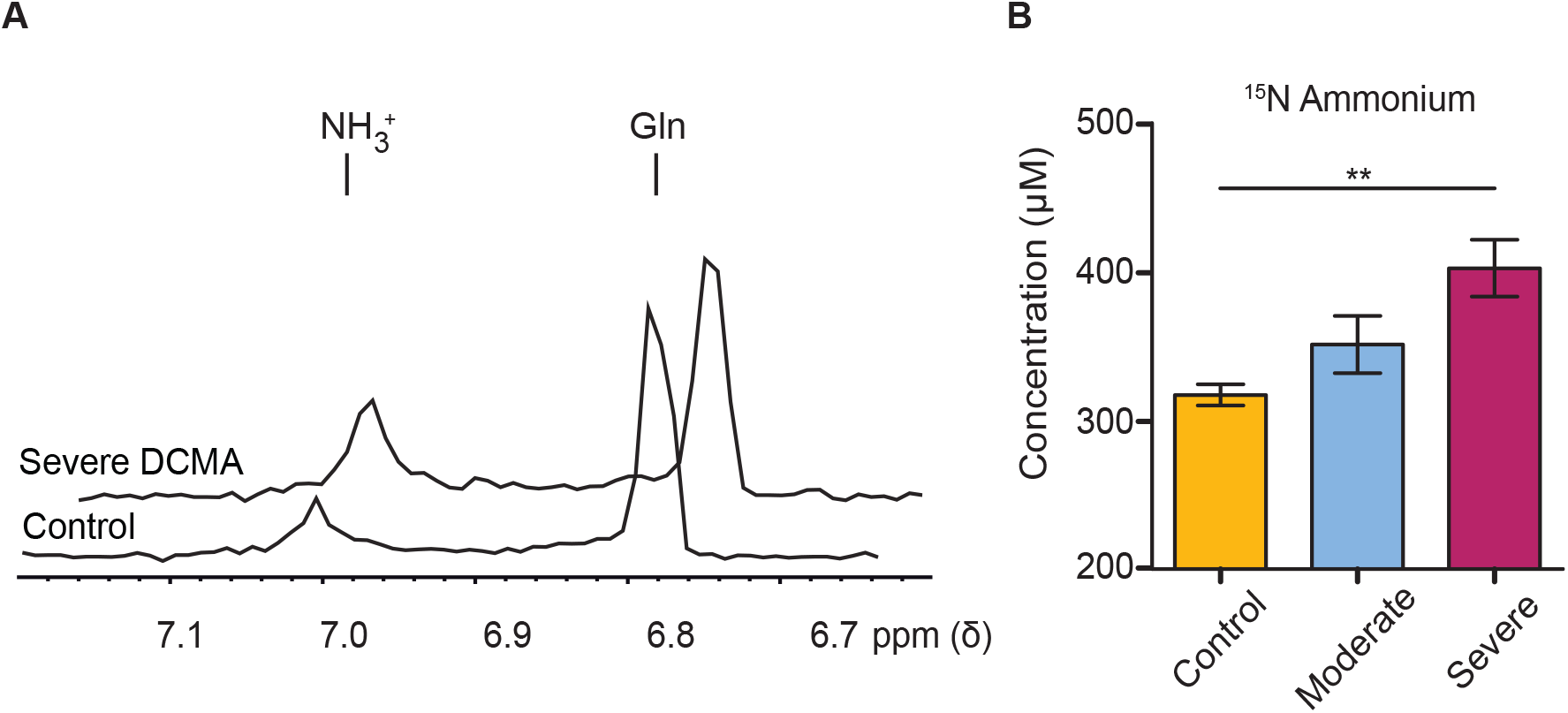
Ammonium secretion in DCMA cell lines. **(A)** ^1^H-^15^N HSQC NMR spectra of control and severe DCMA medium samples. The ammonium peak is highlighted at 7.01 ppm and glutamine at 6.81 ppm. **(B)** ^15^N-labelled ammonium concentration in fibroblast medium after 24 hours incubation. Dermal fibroblasts from non-DCMA healthy controls and DCMA patients with mild and severe cardiac dysfunction were incubated with [5-^13^C 2-^15^N] stable isotope-labelled glutamine in DMEM. Data are the mean ± SEM of three biological replicates per condition, with two technical replicates each. ** p < 0.005 using Student’s two-sided t-test.

### Predicting DCMA disease progression via glutamine recycling

Our data show that glutamine recycling, as measured by [5-^13^C, 1-^15^N] and [5-^13^C] glutamine levels, is one of the most significant metabolic abnormalities found in DCMA cell lines. Given that these phenotypes appeared to correlate with DCMA severity in our initial data set, we hypothesized that glutamine recycling may serve as a potential predictive tool to identify patients with severe heart disease. To test this hypothesis, we analyzed glutamine utilization in an independent cohort of DCMA cell lines (n=9) in a blinded manner. None of the metabolic profiles of the blinded cohort were accessible to the study team or had been analyzed prior to this trial. Glutamine recycling phenotypes were quantified using our MS analytical approach and results were standardized relative to the mild DCMA cell lines used in our training data sets. The clinical severity of cell lines in our blinded cohorts was then predicted based on the distribution of the glutamine recycling phenotypes observed in the mild reference cell lines. The boundary between severe and moderate classifications was defined as 2 standard deviations below the normal distribution observed in the reference cell lines. Clinical information on patients was maintained blinded until predictions were made. No modifications, additions, or exclusions were made to the validation data set and the validation data had never been previously used to assess or refine the model being tested.

Our metabolomics-based diagnostic tool correctly differentiated severe from mild DCMA cases with only one incorrect patient classification (Figure 5). Both [5-^13^C 1-^15^N] glutamine and [5-^13^C] glutamine levels correctly separated the two classes. Overall, the positive predictive value (PPV) for glutamine recycling was 1.0 and the negative predictive value (NPV) was 0.83.

**Figure 5.**
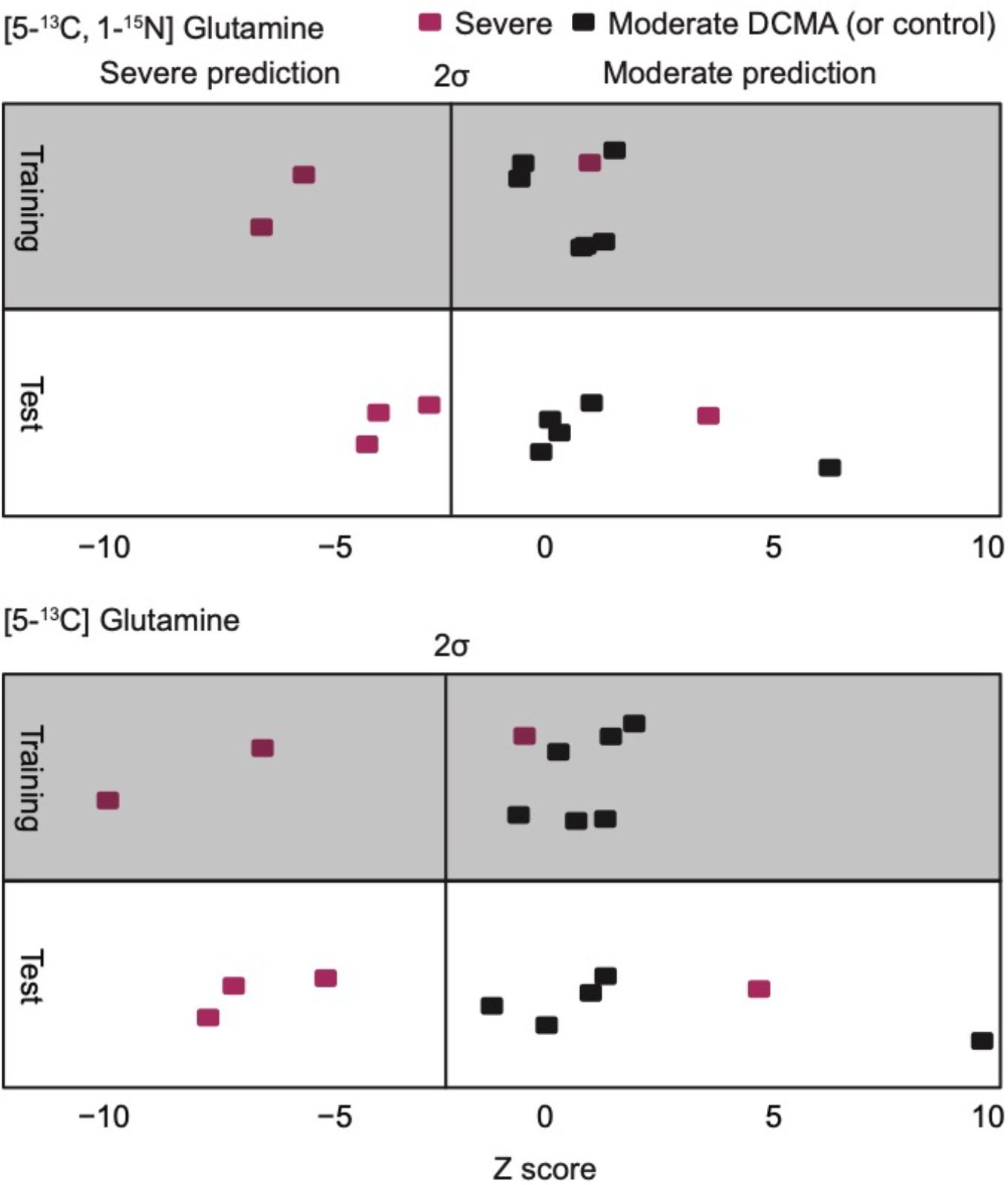
Prediction of DCMA severity from glutamine recycling. **(A)** Results from both the training and validation (test) cohorts after incubation with 5-^13^C 1-^15^N glutamine. **(B)** Results from both the training and validation cohorts after incubation with 5-^13^C glutamine. Each point represents the mean of three technical replicates of glutamine labelling levels observed in cultured cell lines after 48 hours incubation with 5-^13^C 1-^15^N glutamine or 5-^13^C glutamine. Metabolite concentrations are expressed as Z-scores relative to mild DCMA cell lines in our training cohort. Mean concentrations of the diagnostic isotopomers observed from each cell line were subtracted from the mean concentrations observed in the mild DCMA cell lines and divided by the standard deviation of concentrations observed in the mild DCMA cell lines. A threshold of 2 standard deviations was set as the default classifier for segregating severe versus mild DCMA cases. Red squares represent severe DCMA cases and black squares represent healthy controls or DCMA patients with mild disease.

## Discussion

Diagnostic tools that can predict the clinical trajectory of DCMA could have a significant impact on the management of this debilitating and frequently lethal disease. DCMA cardiac clinical phenotypes range from mild cardiac dysfunction to death before 2 years of age due to severe heart failure that is unresponsive to standard medical therapy. Tools for identifying at-risk patients would enable effective medical interventions to be employed before the disease becomes life-threatening. The primary objective of this study was to conduct a detailed analysis of the metabolic consequences of DCMA and identify metabolic phenotypes that are predictive of clinical severity. We report a novel glutamine catabolism phenotype that successfully differentiated severe from mild cases in a blinded cohort of samples. These findings highlight a previously unrecognized and significant metabolic abnormality in DCMA that may provide a diagnostic tool for predicting the clinical trajectory of this potentially life-threatening disease.

The biological rationale for this DCMA-linked perturbation in glutamine metabolism is unclear. However, one possibility is that *DNAJC19* mutations may affect the mitochondria’s ability to import glutamate, and thus elicits elevated glutamine metabolic fluxes. These glutamine-dependent reactions may serve as a compensatory mechanism to support overall TCA activity. This interpretation is supported by the observation that, collectively, only 33.6% (335 µM) of the glutamine brought into the cells was secreted as glutamate and glutamine, with the remainder entering TCA metabolism and biosynthetic reactions. Alternatively, liberating ammonium may simply provide a counter ion for facilitating carbon transport that compensates for some, as of yet unidentified transport defect caused by DCMA. Importantly, a recent study *(15)* revealed that DNAJC19 interacts with numerous mitochondrial transport proteins (Figure S3 and Table S2). Mutations in DNAJC19 could thus interfere with these interactions and alter the permeability of the mitochondrial membrane, making the glutamine metabolic phenotype in DCMA cells a transport-driven phenomenon.

Alterations in the mitochondrial membrane may also explain another one of our core metabolic findings: the elevated anaerobic catabolism of glucose. Most of the ATP needed by fibroblasts is supplied by the anaerobic catabolism of glucose. Our data support this mechanism and show that DCMA cell lines have slightly elevated glycolytic flux. These elevated fluxes may be partially attributable to the glutamine uptake and utilization, which require active transport mechanisms to move glutamine from the medium to the cytosol, then cytosol to the mitochondrion *(16)*. Moreover, the previously reported mitochondrial fragmentation may have more global energetic impacts on DCMA cell lines and thus require elevated glycolytic activities. These data may also help explain why we have seen that the administration of glucose to DCMA patients is therapeutically beneficial, particularly during fasting or physiological stress.

Another interesting observation was the uniform carbon partitioning we observed in central carbon metabolism. Increased glutamine and glucose utilization would normally be interpreted as evidence for altered intracellular carbon utilization. However, our intracellular labelling data collected during steady-state conditions clearly indicates that the overall fractions of glutamine versus alternative nutrient sources in the mitochondria of DCMA patients remains constant, despite the elevated glutamine utilization. Our data shows that much of this glutamine is secreted as glutamate and ammonium, which provides a molecular interpretation for this phenotype. The glutaminase enzyme necessary for this conversion is localized to the mitochondrial matrix and thus requires transport reactions both to access the glutamine and secrete the resulting glutamate and ammonium. These data along, with the lack of any detectable changes in intracellular fluxes, the apparent independence of the phenotype from oxygen tension, and the presence of exogenous glutamate collectively suggest that increased glutamine metabolic flux in DCMA cells is not secondary to energy or biomass production needs but rather a direct result of the *DNAJC19* mutation.

One of the most exciting outcomes of this study is the fact that our novel glutamine phenotype appears to track with the clinical classification of the disease. Although we report here a range of metabolic phenotypes, glutamine recycling appears to be the only phenotype we observed that can differentiate severe cases from those with mild disease. In contrast, the most characteristic DCMA-linked metabolic phenotype, the production of 3-MGA and 3-MGC, was not observed in patient-derived fibroblasts. This observation is likely attributable to differential metabolism amongst various cell types.

Although the rare nature of DCMA precludes a large-scale blinded trial, our work represents the largest number of patient-derived cells studied to date. Although the cell culture-based fibroblast system used here would be an inconvenient clinical assay for logistical reasons, a similar assay could potentially be developed using peripheral blood mononuclear cells, which would be more easily accessible and permit rapid diagnostic testing. Moreover, this diagnostic strategy based upon abnormal glutamine catabolism may have wider applicability to other mitochondrial metabolic disorders.

## Materials and Methods

### Patient fibroblasts

This study was approved by the Conjoint Health Research Ethics Board at the University of Calgary. Dermal fibroblast cell lines collected from Hutterite DCMA patients during clinical investigation were stored in liquid nitrogen in the Biochemical Genetics laboratory facility at the Alberta Children’s Hospital. Commercially-available control, non-Hutterite fibroblast cell lines were obtained from ThermoFisher Scientific and the Coriell Institute for Medical Research. Fibroblasts were maintained in Dulbecco’s Modified Eagle Media (DMEM) supplemented with 2 mM glutamine, 100 U/mL penicillin/streptomycin, and 10% fetal bovine serum (FBS) (ThermoFisher Scientific) and grown in T75 flasks in a humidified incubator at 37°C and 5% CO2 with medium changes every 4-6 days. For metabolomics experiments, maintenance DMEM was changed to stable isotope-labelled DMEM. This DMEM contained 2 mM [5-^13^C 2-^15^N] labelled glutamine (Cambridge Isotope Laboratories, Inc.), 5.5 mM glucose, and 10% dialyzed FBS (ThermoFisher Scientific). Experimental glucose concentration was lowered from 25 to 5.5 mM to better represent *in vivo* glucose levels *(9)*.

Three fibroblast cell lines were derived from DCMA patients with severe cardiac dysfunction (S1-S3), three from patients with mild cardiac dysfunction (M1-M3), and three unrelated control fibroblast cell lines (C1-C3) (Table 1). Two control cell lines C1 and C2 (human dermal fibroblast adult, or hDFa lines) were isolated from healthy adults, and one (C3) was isolated from a healthy control, age-matched to all severe DCMA patient cell lines. For our validation study, nine fibroblast cell lines obtained from DCMA patients with a range of clinical presentations were used, and the clinical data was revealed only after predictions were made based on the assay. No outliers were excluded from the study. Biological replicates are defined as fibroblasts derived from different patients. Technical replicates are experimental replicates derived from different flasks or wells of the same biological replicate.

For metabolomics experiments on media, fibroblasts were seeded in tissue culture-treated 24-well plates at a seeding density of 5.0 × 10^4^ live cells and were allowed to grow to confluency for 24 hours. At t=0 h, the unlabeled DMEM was aspirated off, cells were washed with 1 mL PBS, and 0.5 mL of the isotope-labelled DMEM was added. Extracellular metabolite extractions were taken at t=48 h, diluted 20-fold in 50% methanol, and stored at −80°C until MS analysis.

### Intracellular carbon partitioning of TCA cycle metabolites and extracellular ammonium assay

A total of 1.0 × 10^6^ live fibroblast cells were seeded into T75 flasks and allowed to grow to confluency for 24 hours. This experiment was performed once with three biological replicates per condition (n=3), and two technical replicates per biological replicate. Fibroblasts were incubated in [5-^13^C] glutamine DMEM for 24 hours. At t=24, medium was collected for ammonium quantification by NMR and intracellular metabolites were extracted with 7 mL cold 90% methanol for analysis by LC-MS. These extracts were 14x concentrated and resuspended in 50% methanol for MS analysis. Carbon partitioning percentages for intracellular TCA cycle metabolites were calculated using the following equation:

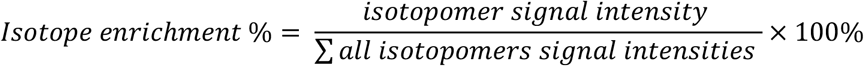

### LC-MS Analysis

All metabolomics analyses were conducted at the Calgary Metabolomics Research Facility (CMRF). Metabolite extracts were analyzed using a ThermoFisher Scientific Q Exactive Basic Orbitrap Mass Analyzer (ThermoFisher Scientific) in negative mode. Liquid chromatography (LC) separation of metabolites was performed on a ThermoFisher Scientific Vanquish ultra high-performance LC (UHPLC) platform. Intracellular metabolites were analyzed using a reverse phase ion pairing (RPIP) strategy adapted from previous methods *(17)* on an Agilent Zorbax RRHD SB-C18 1.5 µM threaded column. The following modifications were made: samples were ran for 15 minutes at a flow rate of 600 µL/min on a four stage linear gradient. Solvent A was titrated to pH 7.5, solvent B was acetonitrile, and injection volume was 2 µL. The gradient was 0 min, 0% B; 1 min, 0% B; 2 min, 25 % B; 3.5 min, 25% B; 5 min, 55% B; 6.5 min, 55% B; 8.5 min, 100% B; 10 min, 100% B; 11 min, 0% B. Medium extracts were analyzed using a hydrophilic interaction LC (HILIC)-based approach on a ThermoFisher Synchronis HILIC column (ThermoFisher Scientific). The HILIC method ran for five minutes at a flow rate of 600 µL/min on a three-stage gradient with solvent A as acetonitrile with 0.1% formic acid and solvent B as 20 mM ammonium formate at pH 3.0. The gradient was 0 min, 0% B; 0.5 min, 0% B; 1.75 min, 20 % B; 3 min, 95% B; 3.5 min, 95% B; 4 min, 0% B. The flow rate was 600 µL/min and total run time was 5 minutes. Matrix-matched external calibration curves of metabolite standards were run alongside biological extracts for quantification of extracellular metabolites. Data were acquired in negative mode with a scan range of 70-1000 m/z, resolution of 140,000, automatic gain control target of 1e6 (RPIP) and 3e6 (HILIC), and maximum injection time of 200 ms.

### NMR Analysis

Extracellular ^15^N-labelled ammonium was detected and quantified using proton 1D ^1^H ^15^N heteronuclear single quantum coherence (HSQC) NMR spectroscopy. Spent medium, collected after 24 h incubation in 10% D2O, was titrated to pH 3 (+/- 0.03). Spectra were then acquired on 600 MHz Bruner NMR spectrometer at 298 K using pulse program hsqcetfpf3gpsi, 64 scans, spectral width of 9615.38 Hz and acquisition time of 0.213 seconds. A matrix-matched external ^15^N ammonium chloride calibration curve (Cambridge Isotope Laboratories) was used to confirm compound IDs and calibrate the quantification of the ammonium following established methods *(18)*. The accurate linear range for quantification was determined to be 150 µM to 750 µM and was used to quantify *in vitro* samples. All NMR spectra were processed using TopSpin version 4.0.7 and analyzed using rNMR.

### Metabolomics-based clinical diagnostic assay

A blinded cohort of fibroblasts (Table 2) was obtained from the Biochemical Genetics Laboratory at the Alberta Children’s Hospital (as described above) and the clinical severity of these patients was kept blinded. These fibroblasts were cultured and analyzed using the previously reported methods. Fibroblasts were incubated with DMEM containing dual stable isotope (^13^C and ^15^N)-labelled glutamine. The training set experiment was performed four times with three biological replicates per condition (n=3), and three technical replicates per biological replicate. The test set experiment was performed one time with three technical replicates per blinded cell line.

### Statistics

Significance was determined using unpaired Student’s two-sided t-tests (data were found to be normally distributed using a D’Agostino and Pearson omnibus normality distribution test, otherwise a Gaussian Approximation was used). P values < 0.05 were considered statistically significant. Data are presented as mean ± standard error of the mean (SEM). P values reported in this manuscript are from unpaired two tailed equal variance t-tests computed in Prism7 (GraphPad). Clinical severity predictions were computed from Z-scores wherein the mean population was defined as the mild DCMA cases and the Z-score was based on secreted glutamine biomarker concentrations of the mild DCMA population. The threshold for severe disease was two standard deviations from the mild DCMA population mean (Z-score = ± 2). Predictions were confirmed by reference to the clinical data available (Table 2).

## Supplementary Figures

**Figure S1.**
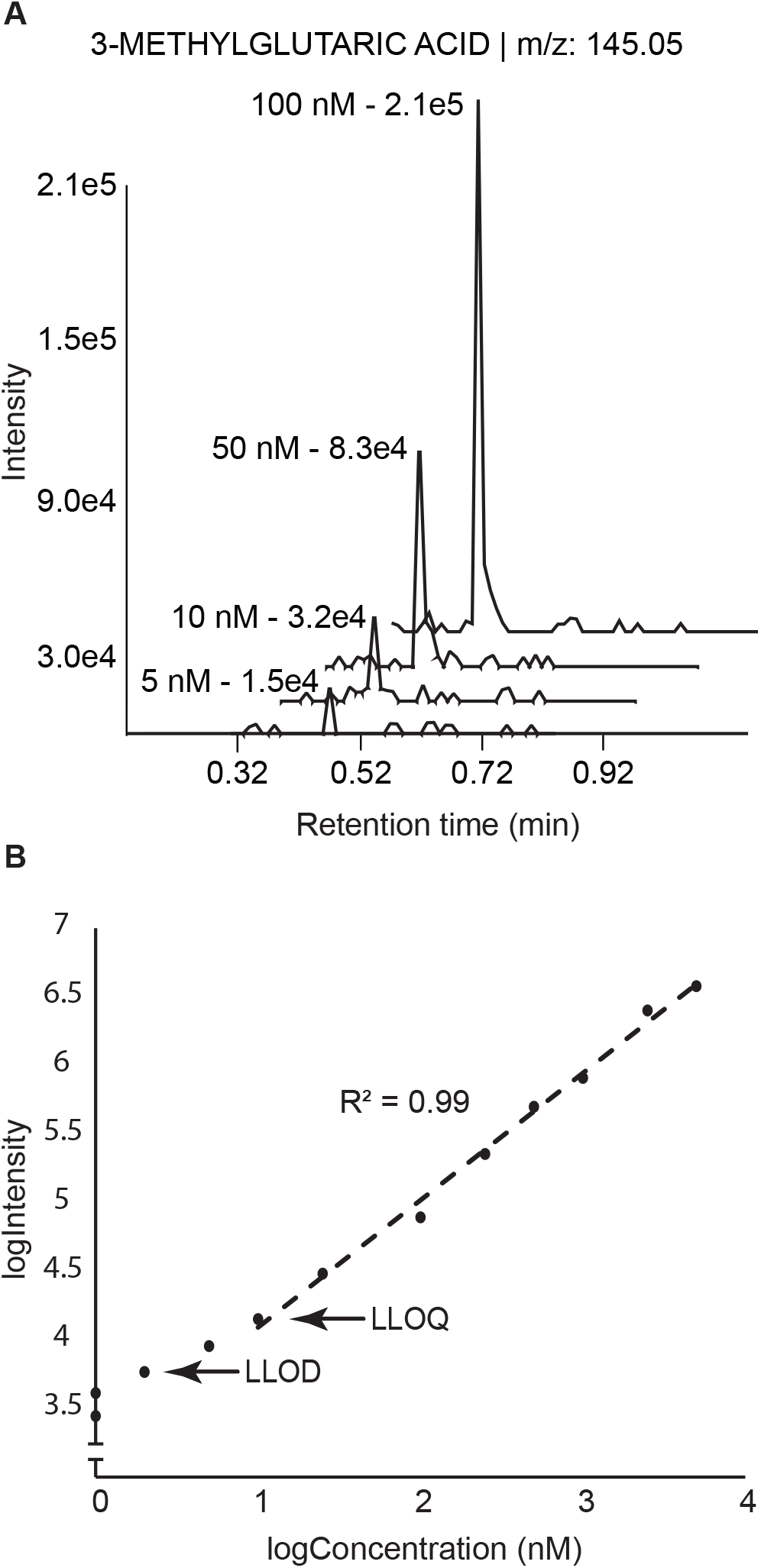
Nanomolar 3-MGA quantification and detection. The extracted ion chromatogram (EIC, top) shows peaks of 3-MGA standard samples. m/z represents mass over charge ratio of metabolite. Log regression curve shows the limit of quantification (LOQ, 100 nM) and limit of detection (LOD, 2 nM) for 3-MGA.

**Table S1.** Live cell counts for fibroblast cell lines.

|  | Live cell count<br>(cells/mL) | Viability (%) | A (cells/mL) |
| --- | --- | --- | --- |
| C1 | 1.00E+06 | 90 | 880,000 |
|  | 7.60E+05 | 88 |  |
| C2 | 600000 | 85 | 600,000 |
|  | 600000 | 81 |  |
| C3 | 880000 | 87 | 900,000 |
|  | 920000 | 83 |  |
| M1 | 940000 | 89 | 1,120,000 |
|  | 1300000 | 86 |  |
| M2 | 450000 | 73 | 485,000 |
|  | 520000 | 83 |  |
| M3 | 660000 | 87 | 700,000 |
|  | 740000 | 82 |  |
| S1 | 300000 | 82 | 300,000 |
|  | 300000 | 86 |  |
| S2 | 470000 | 79 | 470,000 |
|  | 470000 | 80 |  |
| S3 | 930000 | 70 | 900,000 |
|  | 870000 | 67 |  |
| U1 | 1.10E+06 | 95 | 1.01E+06 |
|  | 9.20E+05 | 94 |  |
| U2 | 1.20E+06 | 82 | 1.20E+06 |
|  | 1.20E+06 | 74 |  |
| U3 | 1.00E+06 | 83 | 1.00E+06 |
|  | 1.00E+06 | 85 |  |
| U4 | 8.70E+05 | 92 | 9.10E+05 |
|  | 9.50E+05 | 90 |  |
| U5 | 1.20E+06 | 88 | 1.15E+06 |
|  | 1.10E+06 | 87 |  |
| U6 | 9.30E+05 | 89 | 9.20E+05 |
|  | 9.10E+05 | 95 |  |
| U7 | 8.40E+05 | 76 | 8.65E+05 |
|  | 8.90E+05 | 78 |  |
| U8 | 1.00E+06 | 91 | 1.10E+06 |
|  | 1.20E+06 | 89 |  |
| U9 | 1.30E+06 | 91 | 1.20E+06 |
|  | 1.10E+06 | 86 |  |

**Figure S2.**
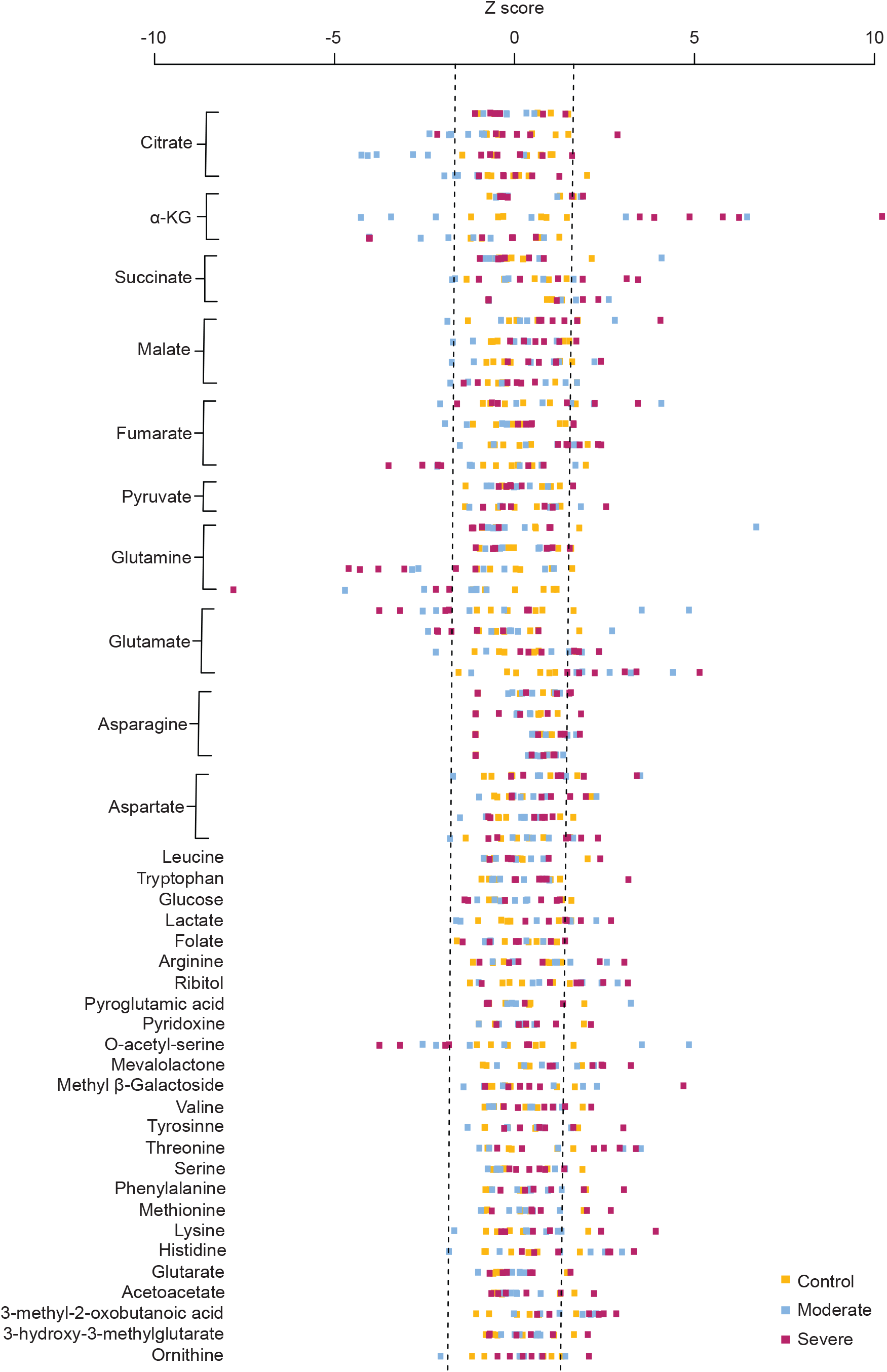
Z-scores of intracellular metabolites of DCMA fibroblasts against control population fibroblasts. Fibroblasts from controls and DCMA patients with mild and severe cardiac dysfunction were provided [5-^13^C 2-^15^N] stable isotope-labelled glutamine in DMEM and incubated for 72 + 24 hours. Z-scores were generated using signal intensities of intracellular metabolites with various labelling partitions.

**Figure S3.**
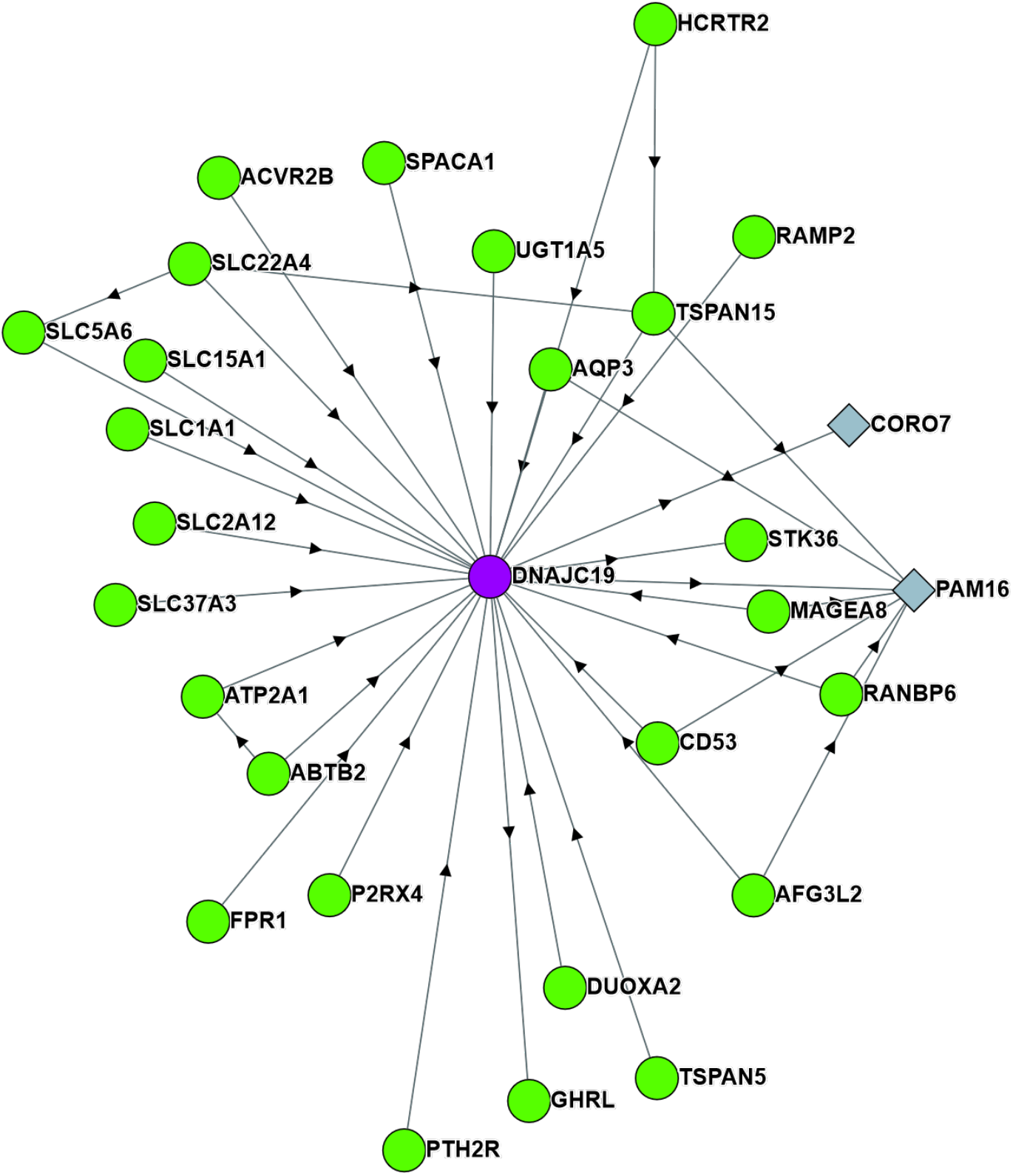
DNAJC19 protein-protein interactome. Interacting partners are represented by green circles and blue diamonds.

**Table S2.** DNAJC19 interacting transporter proteins and their known or proposed functions. Interacting partners are ordered from highest to lowest probability of interaction.

| **Protein** | **Proposed cellular function** |
| --- | --- |
| SLC5A6<br>(Solute carrier family 5 member 6) | Transports pantothenate (vitamin B5), biotin and lipoate in the presence of sodium. |
| SLC1A1<br>(Solute carrier family 1 member 1) | High-affinity glutamate transporter, also transports aspartate |
| SLC22A4<br>(Solute carrier family 22 member 4) | Related to cation (sodium) transport, transport by this protein is ATP dependent |
| SLC37A3<br>(Solute carrier family 37 member 3) | Glucose-6-P or glycerol-3-P transmembrane transporter |
| SLC15A1<br>(Solute carrier family 15 member 1) | Protein uptake in small intestine epithelium |
| SLC2A12<br>(Solute carrier family 2 member 12) | Glucose transporter, GLUT12, homology to insulin-receptive GLUT4 |

## References

1. K. M. Davey, J. S. Parboosingh, D. R. McLeod, A. Chan, R. Casey, P. Ferreira, F. F. Snyder, P. J. Bridge, F. P. Bernier, Mutation of DNAJC19, a human homologue of yeast inner mitochondrial membrane co-chaperones, causes DCMA syndrome, a novel autosomal recessive Barth syndrome-like condition, J. Med. Genet. 43, 385–93 (2006).

2. L. Rohani, G. Meng, P. Machiraju, S. Liu, J. Wu, I. Kovalchuk, I. Lewis, T. Shutt, A. Khan, D. Rancourt, S. Greenway, Modeling the Dilated Cardiomyopathy with Ataxia Syndrome (DCMA), A Pediatric Mitochondrial Cardoimyopathy, Using Cardiomyocytes Derived from Induced Pluripotent Stem Cells, Can. J. Cardiol. 33, S163–S164 (2017).

3. A. Al Teneiji, K. Siriwardena, K. George, S. Mital, S. Mercimek-Mahmutoglu, Progressive Cerebellar Atrophy and a Novel Homozygous Pathogenic DNAJC19 Variant as a Cause of Dilated Cardiomyopathy Ataxia Syndrome, Pediatr. Neurol. 62, 58–61 (2016).

4. T. Ojala, P. Polinati, T. Manninen, A. Hiippala, J. Rajantie, R. Karikoski, A. Suomalainen, T. Tyni, New mutation of mitochondrial DNAJC19 causing dilated and noncompaction cardiomyopathy, anemia, ataxia, and male genital anomalies, Pediatr. Res. 72, 432–437 (2012).

5. A. Chowdhury, A. Aich, G. Jain, K. Wozny, C. Lüchtenborg, M. Hartmann, O. Bernhard, M. Balleiniger, E. A. Alfar, A. Zieseniss, K. Toischer, K. Guan, S. O. Rizzoli, B. Brügger, A. Fischer, D. M. Katschinski, P. Rehling, J. Dudek, Defective Mitochondrial Cardiolipin Remodeling Dampens HIF-1α Expression in Hypoxia, Cell Rep. 25, 561-570.e6 (2018).

6. P. Machiraju, X. Wang, R. Sabouny, J. Huang, T. Zhao, F. Iqbal, M. King, D. Prasher, A. Lodha, A. Ravandi, B. Argiropoulos, D. Sinasac, A. Khan, T. Shutt, S. C. Greenway, SS-31 Reverses Mitochondrial Fragmentation in Fibroblasts from Patients with DCMA, a Mitochondrial Cardiomyopathy, bioRxiv, 672857 (2019).

7. S. B. Wortmann, L. A. Kluijtmans, U. F. H. Engelke, R. A. Wevers, E. Morava, The 3-methylglutaconic acidurias: what’s new?, J. Inherit. Metab. Dis. 35, 13–22 (2012).

8. R. Richter-Dennerlein, A. Korwitz, M. Haag, T. Tatsuta, S. Dargazanli, M. Baker, T. Decker, T. Lamkemeyer, E. I. Rugarli, T. Langer, DNAJC19, a Mitochondrial Cochaperone Associated with Cardiomyopathy, Forms a Complex with Prohibitins to Regulate Cardiolipin Remodeling, Cell Metab. 20, 158–171 (2014).

9. J. M. S. Lemons, H. A. Coller, X. J. Feng, B. D. Bennett, A. Legesse-Miller, E. L. Johnson, I. Raitman, E. A. Pollina, H. A. Rabitz, J. D. Rabinowitz, Quiescent fibroblasts exhibit high metabolic activity, PLoS Biol. 8 (2010).

10. H. Ke, I. A. Lewis, J. M. Morrisey, K. J. McLean, S. M. Ganesan, H. J. Painter, M. W. Mather, M. Jacobs-Lorena, M. Llinás, A. B. Vaidya, Genetic investigation of tricarboxylic acid metabolism during the plasmodium falciparum life cycle, Cell Rep. 11, 164–174 (2015).

11. R. I. Kelley, Quantification of 3-methylglutaconic acid in urine, plasma, and amniotic fluid by isotope-dilution gas chromatography/mass spectrometry, Clin. Chim. Acta 220, 157–164 (1993).

12. Y. D. Bhutia, V. Ganapathy, Glutamine transporters in mammalian cells and their functions in physiology and cancerBiochim. Biophys. Acta - Mol. Cell Res. (2016), doi:10.1016/j.bbamcr.2015.12.017.

13. I. A. Lewis, S. C. Schommer, B. Hodis, K. A. Robb, M. Tonelli, W. M. Westler, M. R. Sussman, J. L. Markley, Method for determining molar concentrations of metabolites in complex solutions from two-dimensional 1H-13C NMR spectra, Anal. Chem. 79, 9385–9390 (2007).

14. I. A. Lewis, S. C. Schommer, J. L. Markley, rNMR: Open source software for identifying and quantifying metabolites in NMR spectra, Magn. Reson. Chem. 47, S123 (2009).

15. E. L. Huttlin, R. J. Bruckner, J. A. Paulo, J. R. Cannon, L. Ting, K. Baltier, G. Colby, F. Gebreab, M. P. Gygi, H. Parzen, J. Szpyt, S. Tam, G. Zarraga, L. Pontano-Vaites, S. Swarup, A. E. White, D. K. Schweppe, R. Rad, B. K. Erickson, R. A. Obar, K. G. Guruharsha, K. Li, S. Artavanis-Tsakonas, S. P. Gygi, J. Wade Harper, Architecture of the human interactome defines protein communities and disease networks, Nature 545, 505–509 (2017).

16. M. Scalise, L. Pochini, M. Galluccio, C. Indiveri, Glutamine transport. From energy supply to sensing and beyond, Biochim. Biophys. Acta - Bioenerg. 1857, 1147–1157 (2016).

17. W. Lu, M. F. Clasquin, E. Melamud, D. Amador-Noguez, A. A. Caudy, J. D. Rabinowitz, Metabolomic analysis via reversed-phase ion-pairing liquid chromatography coupled to a stand alone orbitrap mass spectrometer., Anal. Chem. 82, 3212–21 (2010).

18. I. A. Lewis, R. H. Karsten, M. E. Norton, M. Tonelli, W. M. Westler, J. L. Markley, NMR method for measuring carbon-13 isotopic enrichment of metabolites in complex solutions, Anal. Chem. 82, 4558–4563 (2010).

